# Constitutively active STING causes neuroinflammation and degeneration of dopaminergic neurons in mice

**DOI:** 10.1101/2022.02.02.478854

**Authors:** Eva M Szegö, Laura Malz, Nadine Bernhardt, Angela Rösen-Wolff, Björn H. Falkenburger, Hella Luksch

## Abstract

The innate immune system can protect against certain aspects of neurodegenerative diseases, but also contribute to disease progression. Stimulator of interferon genes (STING) is activated after detection of cytoplasmic dsDNA by cGAS (cyclic GMP-AMP synthase) as part of the defense against viral pathogens, activating type I interferon and NF-kB/inflammasome signaling. In order to specifically test the relevance of this pathway for the degeneration of dopaminergic neurons in Parkinson’s disease, we studied a mouse model with heterozygous expression of the constitutively active STING variant N153S.

In adult mice expressing N153S STING, the number of dopaminergic neurons was smaller than in controls, as was the density of dopaminergic axon terminals and the concentration of dopamine in the striatum. We also observed alpha-synuclein pathology and a lower density of synaptic puncta. Neuroinflammation was quantified by staining astroglia and microglia, by measuring mRNAs, proteins and nuclear translocation of transcription factors. Neuroinflammatory markers were already elevated in juvenile mice, thus preceding the degeneration of dopaminergic neurons. Inflammation and neurodegeneration were blunted in mice deficient for signaling by type I interferons or inflammasomes, but not suppressed completely.

Collectively, these findings demonstrate that chronic activation of the STING innate immunity pathway is sufficient to cause degeneration of dopaminergic neurons. This pathway could be targeted therapeutically.

## Introduction

Inflammation contributes significantly to the pathogenesis of neurodegenerative diseases, including Parkinson’s disease (PD) (Harms et al., 2021; Hirsch and Standaert, 2021). Inflammatory serum markers are associated with more severe PD symptoms and with a more rapid progression of cognitive decline (Hall et al., 2018; Mollenhauer et al., 2019). Anti-inflammatory drugs like aspirin have been associated with a lower risk of developing PD in epidemiological studies (Chen et al., 2003). The pathological hallmarks of PD include the degeneration of dopaminergic neurons in the substantia nigra and the cytoplasmic inclusions termed Lewy bodies (Obeso et al., 2017). Aggregates of alpha-synuclein (aSyn), one of the main constituents of Lewy bodies, stimulate monocytes and microglia (Grozdanov et al., 2019) and astroglia (Chou et al., 2021), suggesting that inflammatory changes in PD respond to aSyn pathology and contribute to disease progression (Harms et al., 2021; Hirsch and Standaert, 2021).

The stimulator of interferon genes (STING) responds to cytoplasmic dsDNA as part of the innate immunity defense against viral pathogens (Paul et al., 2021). STING is activated by cyclic GMP-AMP synthase (cGAS), which binds dsDNA and catalyzes synthesis of the second messenger cyclic GMP-AMP (Motwani et al., 2019). This pathway can be activated by viral nucleic acids but also by self-DNA (Chen et al., 2016; Li and Chen, 2018; Motwani et al., 2019). The cGAS-STING pathway has been implicated in the pathogenesis of PD (Beyer et al., 2020), but also Alzheimer’s disease and amyotrophic lateral sclerosis (Chen et al., 2021; Li et al., 2021; Paul et al., 2021). For instance, mice deficient for the PD-associated proteins PINK and parkin fail to degrade damaged mitochondria, accumulate mitochondrial DNA in the cytosol and show a STING-mediated inflammatory phenotype after exhaustive exercise (Sliter et al., 2018).

The two main transcription factors activated by STING are the interferon regulatory factor 3 (IRF3) (Chen et al., 2021) and nuclear factor ’kappa-light-chain-enhancer’ of activated B-cells (NF-κB) (Liu et al., 2014). Consequently, type I interferons (IFN) and pro-inflammatory cytokines are produced, triggering a secondary inflammation through the activation of inflammasomes (Hopfner and Hornung, 2020). Additionally, STING can activate inflammasomes directly (Wang et al., 2020). Inflammasomes are major signaling hubs that activate caspase-1 and control the bioactivity of pro-inflammatory cytokines of the interleukin (IL)-1 family (Gaidt et al., 2017; Schroder and Tschopp, 2010). Activation of inflammasomes formed by the NOD-, LRR- and pyrin domain-containing protein 3 (NLRP3) has been linked to the progression of several neurodegenerative diseases (Heneka et al., 2018). NLRP3 immunoreactivity is increased in mesencephalic neurons of PD patients and NLRP3 variants are associated with the risk to develop PD (von Herrmann et al., 2018). Conversely, inhibiting NLRP3 mitigates degeneration of dopaminergic neurons and aSyn pathology in mouse models (Gordon et al., 2018). NRLP3 has also been associated with Alzheimer’s disease (Heneka et al., 2018), prion disease (Nazmi et al., 2019), and traumatic brain injury (Sen et al., 2020).

Neuroinflammation receives growing attention in neurodegenerative diseases because it represents a promising therapeutic target. Designing such therapies requires, however, to determine the specific effects of individual components of this highly interconnected signaling network, which responds to diverse stimuli and involves many different cell types. Given the role of the STING pathway in PINK/parkin related damage (Sliter et al., 2018), we wanted to learn more about the role of STING for PD pathogenesis and determine whether specific activation of STING is sufficient to cause degeneration of dopaminergic neurons. In order to test this, we used a mouse model with heterozygous expression of the STING genetic variant N153S (Luksch et al., 2019). Constitutively active STING mutants cause an autoinflammatory disease in humans termed STING-associated vasculopathy with onset in infancy (SAVI) (Crow and Casanova, 2014; Liu et al., 2014). SAVI is characterized by systemic inflammation with acral vasculitis, T cell lymphopenia, and interstitial pulmonary disease. Major features of systemic inflammation in SAVI are recapitulated in STING N153S knockin mice (Luksch et al., 2019, Siedel et al., 2020). For simplicity, we refer to these as STING ki mice here and use the term STING WT for the corresponding wild type littermate controls. In this work, we determined the extent of neuroinflammation in the brains of young and adult mice, the extent of dopamine neuron degeneration, and aSyn pathology. Furthermore, we used additional knockout mice to determine which of the known downstream signaling pathways contribute to STING-induced neurodegeneration.

## Materials and Methods

Source of chemicals, antibodies, composition of buffers, equipment and software used in this study are listed in supplemental Table S1.

### Animals

All animal experiments were carried out in accordance with the European Communities Council Directive of November 24, 1986 (86/609/EEC) and approved by the Landesdirektion Dresden, Germany. Mice of both sexes were housed under a 12-hour light and dark cycle with free access to pelleted food and tap water in the Experimental Center, Technische Universität, Dresden, Germany. Heterozygous STING N153S/WT ki mice (STING ki) were previously described (Luksch et al., 2019). STING ki or STING WT mice were crossed to *Ifnar1*^−/−^ mice (a gift from Axel Roers, Dresden, Germany) and *Casp1*^−/−^ mice (a gift from Stefan Winkler, Dresden, Germany).

For all genotypes, five-week-old (from here referred as juvenile) or 20-23-week-old (referred as adult) animals were sacrificed with an overdose of isoflurane (Baxter, Lessines, Belgium). For western blot analysis and for gene expression analysis, brains were rapidly removed from the skull and washed in ice-cold Tris-buffered saline (TBS, pH 7.4). Cortex and striatum were dissected, snap-frozen in liquid nitrogen and stored at −80°C until use. For histology, mice were perfused transcardially with 4 % paraformaldehyde (PFA) in TBS. After post-fixation (4 % PFA, overnight) and cryoprotection (30 % sucrose in TBS), 30 μm-thick coronal brain sections were cut in a cryostat (Leica, Germany).

### Immunofluorescence stainings of mouse brain sections

To quantify the number of dopaminergic neurons in the *substantia nigra* (SN), every fifth section throughout the entire SN was stained for tyrosine hydroxylase (TH) as previously (Szegő et al., 2021). In brief, after blocking (2 % bovine serum albumin, 0,3 % Triton X-100 in TBS; 1 h RT), sections were incubated with the primary antibody in blocking solution (two overnights), followed by the fluorescently labelled secondary antibody (Alexa 488 conjugated donkey anti-sheep, 1:2000, overnight). Sections were counterstained with Hoechst and mounted with Fluoromount-G.

To quantify the density of dopaminergic axon terminals (fibers) and neuroinflammation in the striatum, every sixth section throughout the entire striatum was stained with a cocktail of primary antibodies: TH (Pel Freeze, P40101, 1:1000), Iba1 (Wako, 019-19741, 1:1000) and GFAP (abcam, ab4674, 1:2000). As fluorescently labelled secondary antibodies, Alexa 488 conjugated donkey anti-sheep, Alexa 555 conjugated donkey anti-rabbit, Alexa 647 conjugated donkey anti-chicken were used (1:2000, overnight). Sections were counterstained with Hoechst and mounted with Fluoromount-G.

### Quantification of dopaminergic neuron number, striatal fiber density and gliosis

The number of dopaminergic somata in the SN was determined by supervised manual counting by an investigator blinded to the experimental groups. For each animal, every fifth section throughout the rostro-caudal extent of the SN (2.54 to −3.88 mm posterior to Bregma based on Paxinos and Franklin, 2001) was incorporated into the counting procedure. In each section, z stacks were acquired (step size: 2 μm, 5 slices in total) from both hemispheres with a 20x objective (N.A 0.8, Axio Imager 2, Zeiss). Stacks were stitched to reconstruct the entire SN. After adjusting the threshold and carefully marking the borders of the SN, only TH-positive cell bodies with a visible nucleus in the blue channel were counted by ImageJ (1,53c; Cell Counter plugin).The total number of neurons per SN was estimated by multiplying the counted cell number by five (every fifth section was used for this analysis). For quantification of gliosis, five fluorescent images were acquired from every sixth striatal section stained for GFAP and Iba1 using a 20x objective. After adjusting the threshold and noise removal (Background subtraction, rolling ball radius 50) from the individual images (separately for GFAP and Iba1 channels), the area fraction was determined by ImageJ from ten regions of interest per image. Results were analyzed using a generalized linear mixed model (glm) in RStudio with a hierarchically nested design (expressed as percent area) as previously (Szegő et al., 2021).

From the same striatal sections, the density of the dopaminergic axon terminals (fibers) was determined as described previously (Szegő et al., 2012). In brief, z-stack images were acquired (five planes, 0.5 μm step size, 100x objective, N.A. 1.4; Axio Imager 2, Zeiss). TH-positive fibers were delineated from the maximal intensity projection (ImageJ) after adjusting the threshold, noise removal and binarization, and density was expressed as percent area. Every sixth section per animal, five images per section and ten boxes per image were analyzed in a hierarchically nested design as above.

### Protein analyses

To detect protein changes, cortical and striatal tissue were mechanically lysed in a buffer containing 250 mM sucrose, 50 mM TRIS (pH 7,5), 1 mM EDTA, 5 mM MgCl_2_, 1 % Triton X-100 in the presence of protease and phosphatase inhibitors (MedChem Express) as previously (Szegő et al., 2012). Samples were centrifuged (14000 g, 30 min, 4 °C) and protein concentration in the supernatant was determined with the BCA method (ThermoFisher, Germany). After boiling with 4x Laemmli buffer (1 M Tris pH 6.8, 0.8 % SDS, 40 % glycerol, 5 % β-mercaptoethanol, traces of bromophenol blue, 5 min, 95 °C), 5 μg protein was loaded onto a 4-20 % Tris/glycine SDS gel for western blot analysis. Membranes were fixed in paraformaldehyde (10 min, RT) and blocked with 1 % bovine serum albumin, 0,05 % Tween 20 in TBS. Membranes were incubated first in the presence of antibodies against phosphorylated aSyn and aSyn, (Cell Signaling), then with βIII-tubulin as loading control (overnight, 4 °C). Following washing, membranes were incubated in the presence of horseradish peroxidase-conjugated secondary antibodies (donkey anti-mouse or donkey anti-rabbit). Signal was detected using chemiluminescent substrate and a camera-based system. ImageJ was used to determine the optical density of protein bands and all data were analyzed for each group (n = 5 animals/group) based on 3 independent blots. Optical densities were normalized to the expression of the density of the tubulin loading control of the same sample, and then expressed relative to the WT animals.

### Gene expression analyses

Total RNA was extracted from snap frozen dissected prefrontal cortex tissue by using the RNeasy Mini Kit (Qiagen, Germany) according to the manufacturer’s instructions. cDNA was generated by MMLV reverse transcription (Promega Germany). Quantitative Real Time PCR assays were carried out by using QuantStudio 5 (Thermo Fisher Scientific, Germany) and GoTaq®aPCR Master Mix with SYBR green fluorescence (Promega, Germany). PCR primer sequences were retrieved from the Primer Bank database (Spandidos et al., 2009). Expression of genes was normalized to the expression of the housekeeping genes (*Hprt1*, *Rpl13a*, *Eef2*) and to the STING WT by using the ΔΔCt method. Sequences of primers are listed in supplemental Table S2.

### Statistical analyses

Sample numbers for each analysis are listed in supplemental Table S4. In graphs, markers represent individual animals; lines represent mean and standard deviation (SD) of all animals. Data normality was tested by the Kolmogorov-Smirnov test and graphically by QQ plot (R, version 2.8.0; R Development Core Team 2008). Grubbs test was used to identify outliers. t-test, Mann-Whitney test or two-way ANOVA were performed using GraphPad Prism (Versions 5.01 and 9.0.0). Linear regression was performed using R. For generalized linear mixed-effects model (Szegő et al., 2021), animal, section and image were used as random effects nested within each other (R package: lme4). p values are indicated in the graphs by symbols with * representing p<0.05, ** representing p<0.01, *** representing p<0.001. Exact p values are given in the Figure legends.

## Results

### Neuroinflammation and degeneration of dopaminergic neurons in mice with constitutive STING activation

To characterize the neuronal phenotype in adult STING ki mice, we assessed neuroinflammation and the integrity of the dopaminergic nigrostriatal system. First, we stained striatal sections for Iba1 to determine the activation of microglia. The Iba1 positive area fraction was 9-fold higher in STING ki mice than in STING WT (Figure 1A and E). Activation of astroglia, as determined by GFAP staining, was 26-fold higher in STING ki mice than in STING WT (Figure 1B and F). Thus, STING ki mice exhibit a strong neuroinflammatory phenotype, consistent with increased systemic inflammation (Luksch et al., 2019).

**Figure 1.**
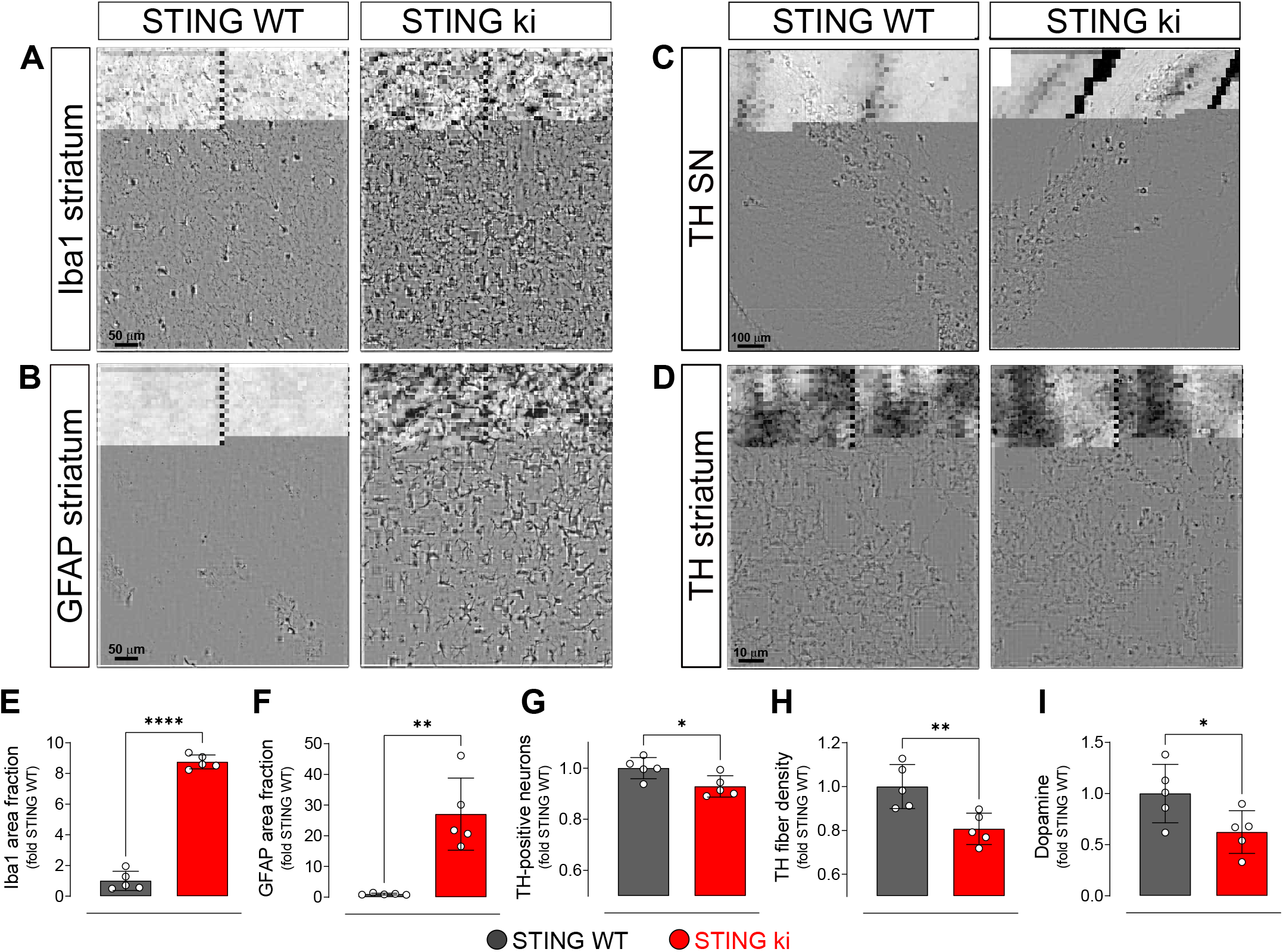
Constitutive STING activation induces neuroinflammation and neurodegeneration in adult mice. (A) Representative images of striatal sections from STING WT and STING ki mice stained for the microglia marker Iba1. Scale bar: 50 μm. (B) Representative images of striatal sections stained for the astroglia marker GFAP. Scale bar: 50 μm. (C) Representative images of midbrain sections containing the *substantia nigra* (SN) from STING WT and STING ki mice (stitched from two microscopy fields) stained for tyrosine hydroxylase (TH). Scale bar: 100 μm. (D) Representative images of striatal sections stained for TH from STING WT and STING ki mice. Scale bar: 10 μm. (E) Area fraction positive for Iba1, normalized to the mean of STING WT mice. Markers represent individual animals (black: STING WT animals, red: STING ki animals). Lines represent mean ± SD. Comparison by t-test (***: p=0,0007). (F) Area fraction positive for GFAP, normalized to the mean of STING WT mice (**: p=0,0011; t-test). (G) Number of TH-positive neurons (*: p=0,0257; t-test). (H) Area fraction positive for TH (**: p=0,0081; t-test). (I) Concentration of dopamine (*: p=0,0448; t-test) in striatal lysates from STING WT and STING ki animals, normalized to the mean concentration in STING WT. Graphs showing quantification of the dopamine metabolites are in figure supplement 1B.

Next, we asked whether the chronic neuroinflammation in the STING ki mice is associated with the degeneration of dopaminergic neurons. Somata of dopaminergic neurons in the *substantia nigra* (SN, Figure 1C) and dopaminergic axon terminals in the striatum (Figure 1D) were identified by staining for tyrosine hydroxylase (TH). The SN of STING ki mice contained significantly fewer TH-positive neurons than the SN of STING WT (Figure 1G and figure supplement 1A). Similarly, the density of dopaminergic axon terminals (fibers) in the striatum was lower in STING ki mice than in STING WT (Figure 1H). Accordingly, the concentration of dopamine in the striatum was lower in STING ki mice than in STING WT mice (Figure 1I). The concentration of dopamine metabolites was higher in STING ki mice (supplemental Figure 1B), suggesting increased dopamine turnover as commonly observed with dopamine depletion. Taken together, these findings demonstrate that the integrity of nigrostriatal dopaminergic neurons is compromised in STING ki mice.

### Neuroinflammation without obvious degeneration of the dopaminergic neurons in juvenile mice with constitutive STING activation

In order to explore whether the compromised integrity of dopaminergic neurons is a consequence of a prolonged neuroinflammation, we next analyzed brain sections of juvenile (5-week-old) STING ki and STING WT mice (Figure 2). Microglia was already significantly activated in juvenile STING ki mice. The area fraction of the Iba1 staining was 2-fold higher in STING ki mice than in STING WT (Figure 2A and E). Similarly, the area fraction of GFAP signal was 14-fold higher in STING ki mice than in STING WT (Figure 2B and F), suggesting activation of astroglia in juvenile STING ki mice. However, the number of TH-positive neurons in the *substantia nigra* was not different between juvenile STING ki mice and STING WT (Figure 2C and G, figure supplement 1C). Similarly, striatal axon terminals (Figure 2D and H) and the concentrations of striatal dopamine and its metabolites (Figure 2I, figure supplement 1D) were not different between STING ki mice and STING WT.

**Figure 2.**
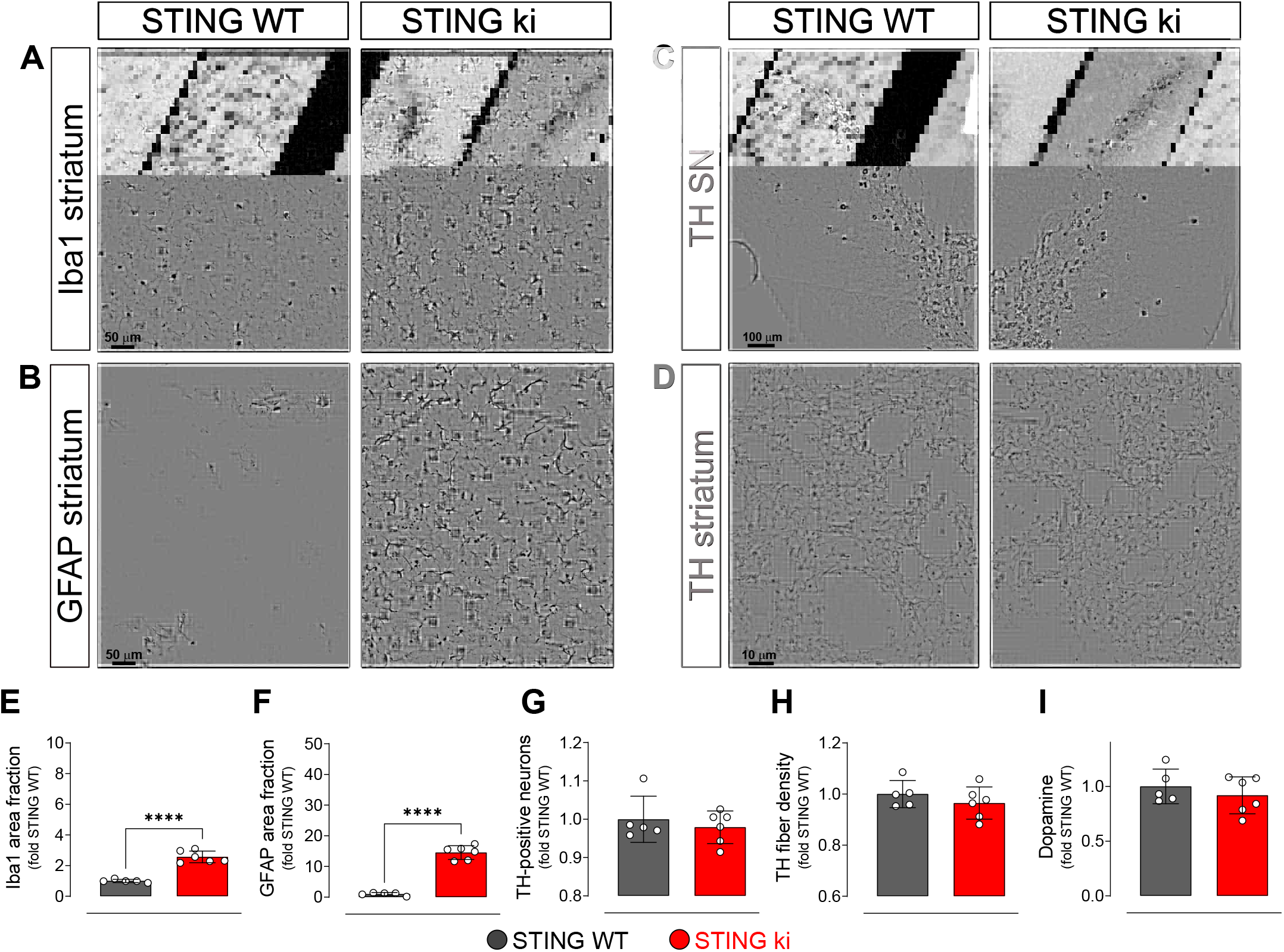
Neuroinflammation without neurodegeneration in juvenile mice with constitutive STING activation. (A) Representative images of striatal sections stained for the microglia marker Iba1 from 5 week-old STING WT and STING ki mice. Scale bar: 50 μm (B) Representative images of striatal sections stained for the astroglia marker GFAP from 5 week-old STING WT and STING ki mice. Scale bar: 50 μm (C) Representative images of midbrain sections containing the *substantia nigra* (SN, stitched from two microscopy fields) stained for tyrosine hydroxylase (TH) from 5 week-old STING WT and STING ki mice. Scale bar: 100 μm (D) Representative images of striatal sections stained for TH from 5 week-old STING WT and STING ki mice. Scale bar: 10 μm. (E) Area fraction positive for Iba1, normalized to the mean of STING WT (***: p=0,0009; t-test). (F) Area fraction positive for GFAP, normalized to the mean of STING WT brains (***: p=0,0007; t-test). (G) Number of TH-positive neurons (mean ± SD; t-test). (H) Area fraction positive for TH (mean ± SD, t-test). (I) Dopamine concentration in striatal lysates from 5 week-old STING WT and STING ki mice, measured by HPLC and normalized to the mean of STING WT (mean ± SD, t-test). Dopamine metabolites are in figure supplement 1D.

Taken together, these findings demonstrate that the compromised integrity of the nigrostriatal system in adult STING ki mice (Figure 1) represents an adult-onset neurodegeneration and not a developmental defect. Given that activation of microglia and astroglia in STING ki mice precedes degeneration of dopaminergic neurons, STING-induced neuroinflammation could contribute to the neurodegeneration.

### aSyn pathology and synaptic defects in the striatum of STING ki mice

To analyze aSyn pathology in the striatum of STING ki mice, we measured the amount of aSyn protein phosphorylated at serine 129 (paSyn), which is considered one of the major pathological forms of aSyn (Anderson et al., 2006; Fujiwara et al., 2002; Samuel et al., 2016). Lysates of adult STING ki mice contained a substantial amount of paSyn (Figure 3A), which was barely detectable in STING WT. The amount of total aSyn was lower in STING ki mice (Figure 3C) and the ratio of paSyn to total aSyn increased (Figure 3D) – as commonly observed in synucleinopathy models and PD patients (Anderson et al., 2006; Chatterjee et al., 2020; Fujiwara et al., 2002; Szegö et al., 2022).

**Figure 3.**
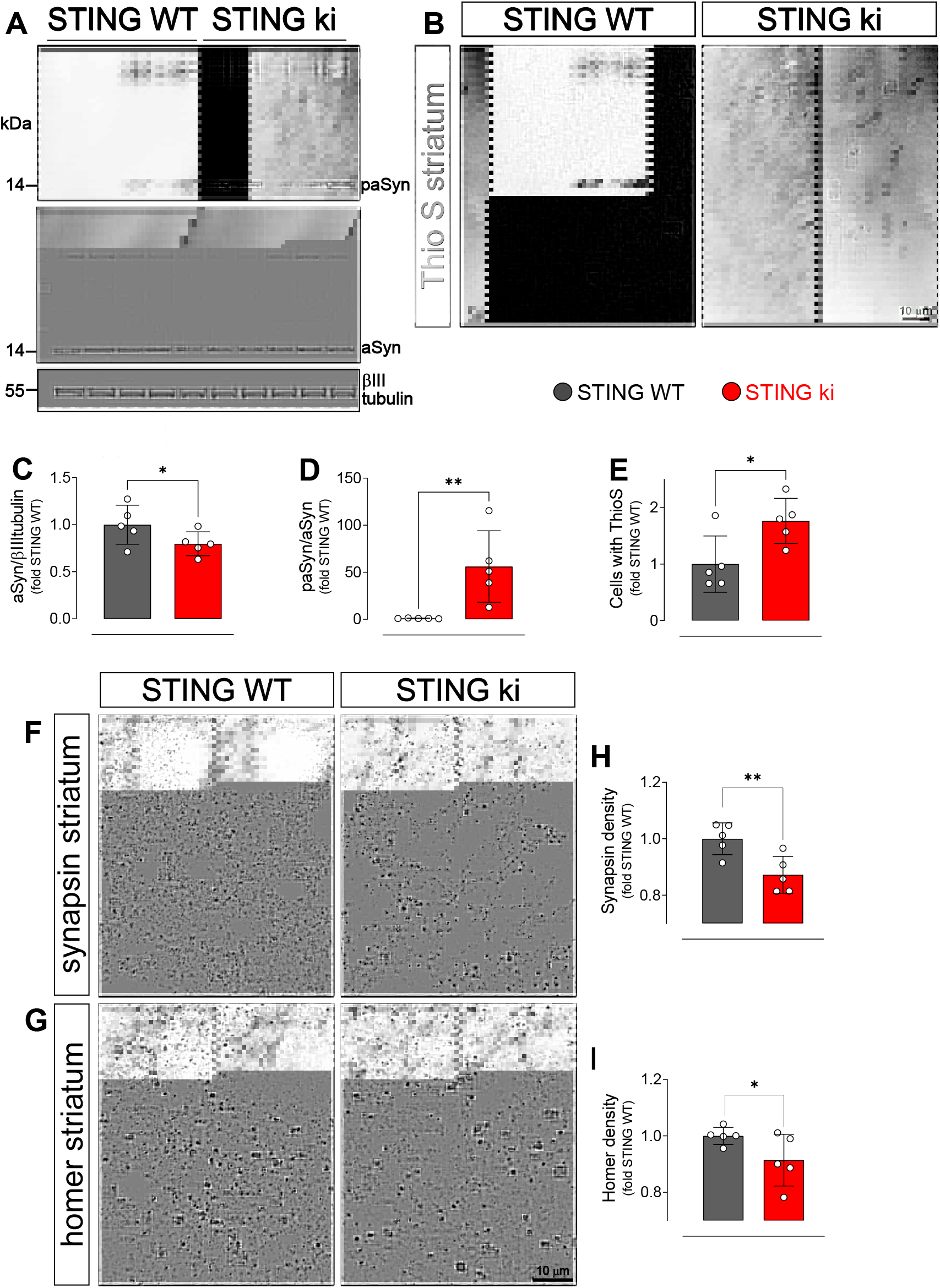
Constitutive STING activation induces alpha-synuclein pathology and synapse loss in adult mice. (A) Representative Western blot images showing phosphorylated alpha-synuclein (S129; paSyn, upper panel), total alpha-synuclein (aSyn, middle panel) and the loading control βIII-tubulin (lower panel). (B) Representative images of striatal sections from 20 week-old STING WT and STING ki mice stained with Thioflavin S. Scale bar: 10 μm (C) Ratio of total alpha-synuclein and loading control, expressed relative to the mean of STING WT (p=0,0491; mean ± SD; t-test). (D) Immunoreactivity to paSyn and aSyn, expressed relative to the mean of STING WT (p=0,0059; mean ± SD; t-test). (E) Number of cells with inclusions positive for Thioflavin S (ThioS) per mm^2^ (*: p=0,0141; t-test). (F-G) Representative images of striatal sections from 20 week-old STING WT and STING ki mice stained for the presynaptic marker synapsin (F) or for the post-synaptic marker homer (G). Scale bar: 10 μm. (H-I) Area fraction positive for synapsin (H, p=0,0053) or homer (I, p=0,0408) (mean ± SD; t-test).

Thioflavin S (ThioS) binds to the characteristic β-sheet conformation of amyloid-containing proteins, including aSyn (Froula et al., 2019; Neumann et al., 2002). The number of cells with ThioS-positive inclusions was higher in the striatum of adult STING ki mice than in STING WT (Figure 3 B and E), consistent with the findings from the aSyn immunoblots.

Since dopamine depletion and aSyn pathology can compromise synaptic integrity, we next quantified the density of synapses in the striatum. Presynaptic puncta were detected by staining against synapsin; post-synaptic puncta were detected by staining against homer (Figure 3F and G). The density of synapsin puncta was 18% lower in adult STING ki than in STING WT (Figure 3H), the density of post-synaptic puncta was 9% lower in STING ki than in STING WT mice (Figure 3I). In summary, we observed aSyn pathology and a reduced density of synapses in the striatum of adult STING ki mice.

### Type I IFN signaling and NF-κB/inflammasome dependent signaling are activated in the brain of STING ki mice

In order to analyze the signaling pathways by which constitutively active STING causes degradation of dopaminergic neurons and aSyn pathology, we examined the expression of selected interferon-stimulated genes (ISGs) by quantitative real time PCR in juvenile and adult STING ki and STING WT mice (Figure 4A-D). In STING WT mice, we observed an age-dependent increase for interferon-induced GTP-binding protein Mx1 (*Mx-1*, 10-fold, p= 0,0000006, comparison not depicted in Figure 4. for clarity), interferon-gamma induced protein 10 kD (*Ip-10*, 22-fold, p= 0,000105) and *Sting1* (3,7-fold, p=0,0008399), consistent with previous findings (Harris et al., 2020).

**Figure 4.**
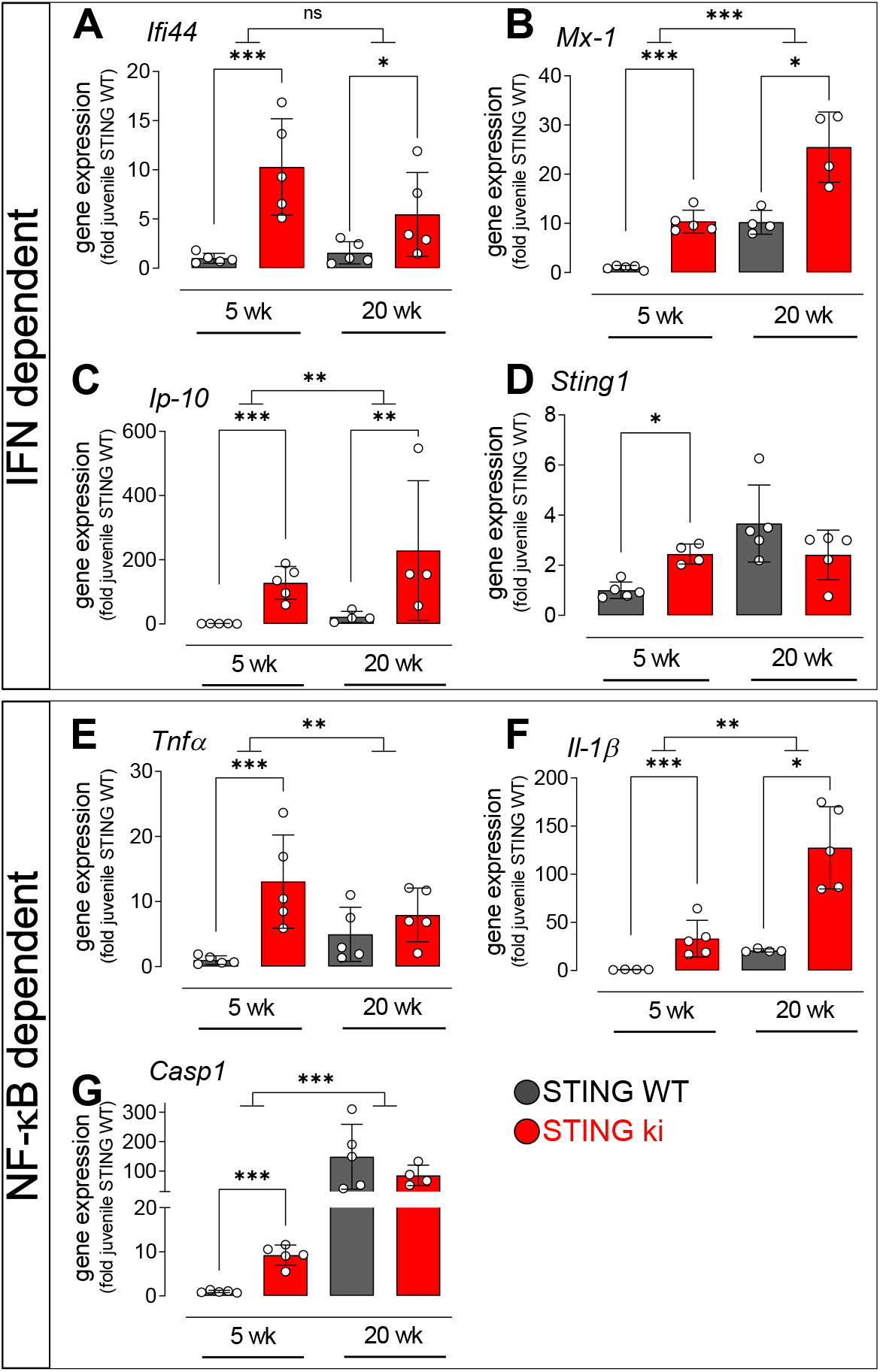
Activation of IFN and NF-κB/inflammasome dependent signaling in juvenile and adult STING ki mice. (A-D) Expression of ISGs in the frontal cortex of STING WT and STING ki mice. (A) *Ifi44* (***: p=0,0002277; *: p= 0,044987), (B) *Mx-1* (***: p=0,0000003; *: p= 0,016835; for comparison between age groups ***: p= 0,000602), (C) *Ip-10* (***: p= 0,000001; **: p= 0,0017215; for comparison between age groups **: p= 0,001483), (D) *Sting1* (*: p= 0,0184042). (E-G) Expression of NF-κB/inflammasome related genes in the frontal cortex of STING WT and STING ki mice. (E) *Tnfα* (***: p= 0,0001448, for comparison between age groups *: p= 0,03952). (F) *Il-1b* (***: p= 0,00005, *: p= 0,0389, for comparison between age groups **: p= 0,00241). (G) *Casp1* (***: p= 0,0000369, for comparison between age groups ***: p= 0,000064). Markers represent individual animals, bars represent mean ± SD. Analysis was two-way ANOVA with Tukey HSD post-hoc test.

In STING ki mice, the expression of ISGs was generally higher than in STING WT, both comparing juvenile mice and comparing adult mice (Figures 4A-D, p-values in Figure legend). The expression of *Sting1* was not significantly different between adult STING ki and STING WT, confirming that the expression of the STING N153S mutant is not changing.

cGAS/STING activation also leads to the activation of NF-κB/inflammasome pathway (Balka et al., 2020; Balka and De Nardo, 2021; Wang et al., 2020). We therefore analyzed induction of the NF-κB/inflammasome pathway by quantitative real time PCR of the downstream mediators, tumor necrosis factor alpha (*Tnfα*), interleukin 1 beta (*Il-1β*), and caspase 1 (*Casp1*) (Figure 4E-G). In STING WT mice, the expression of *Tnfα*, *Il-1β*, *Casp1* increased with age (*Tnfα*: 5-fold, p= 0,0238678; *Il-1β*: 21-fold, p= 0,0000038; *Casp1*: 150-fold, p<0,00001, comparisons not depicted in Figure 4. for clarity), consistent with earlier reports (Mejias et al., 2018). In juvenile STING ki mice, the expression of *Tnfα* was 13-fold higher than in juvenile STING WT (Figure 4E; p= 0,0001448), *Il-1β* was 33-fold higher (Figure 4F, p= 0,00005) and *Casp1* was 9-fold higher (Figure 4G; p= 0,0000369). In adult mice, expression of *Tnfa* and *Casp1* was not different between STING WT and STING ki whereas expression of *Il-1β* was 6-fold higher in adult STING ki than in adult STING WT (Figure 4F, p= 0,0389).

### STAT3 and NF-κB translocate to the nucleus in STING ki mice

Nuclear trafficking is critical for the function of transcription factors such as STAT3 and NF-κB (Balka and De Nardo, 2021; Noguchi et al., 2013). To further investigate the activation of the type I IFN and NF-kB-dependent pathways in juvenile and adult STING ki mice, we quantified the number of nuclei positive for phosphorylated STAT3 (pSTAT3) and NF-κB in the striatum (Figure 5). There were only very few pSTAT3-positive nuclei in juvenile and adult STING WT mice (Figure 5A and C). In juvenile STING ki mice, the average number of pSTAT3-positive nuclei was 4-fold higher than in juvenile STING WT mice (Figure 5C, p= 0,05678, 2-way ANOVA). In adult STING ki mice, the number of pSTAT3-positive nuclei was 15-fold higher than in STING WT (p= 0,00004). Similarly, the number of NF-kB-positive nuclei was 3-fold higher in juvenile STING ki mice than in STING WT (Figure 5B and, p= 0,009). In adult STING ki mice, the number of NF-kB-positive nuclei was 6-fold higher than in STING WT (Figure 5D, p= 0,007).

**Figure 5.**
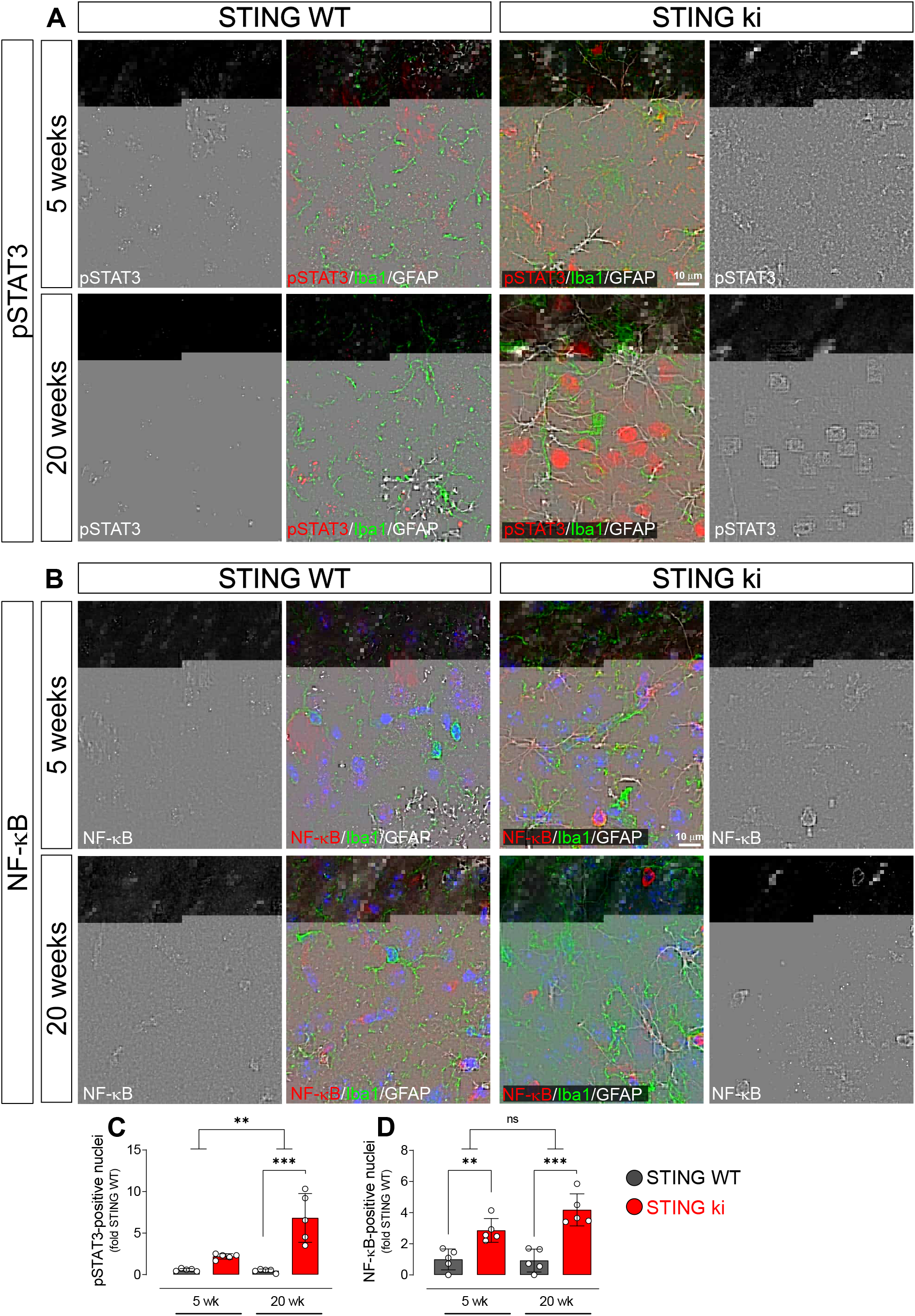
Nuclear translocation of pSTAT3 and NF-κB in the striatum of 5 and 20 week-old STING WT and STING ki mice. (A) Representative images of striatal sections from 5 week-old (upper images) and 20 week-old (lower images) STING WT and STING ki mice stained for Iba1 (green), GFAP (white) and phosphorylated-STAT3 (pSTAT3; red). Images show color coded merged channels (center) and in addition pSTAT3 staining in grayscale (left and right). Scale bar: 10 μm. (B) Representative images of striatal sections from 4 week-old (upper images) and 20 week-old (lower images) STING WT and STING ki mice stained for Iba1 (green), GFAP (white) and NF-κB (red). NF-kB staining is shown in gray in separate images. Scale bar: 10 μm. (C) Number of pSTAT3-positive nuclei/mm^3^ (***: p=0,00004; **: p=0,0025 for the interaction; two-way ANOVA, Bonferroni post-hoc test). (D) Number of NF-kB-positive nuclei/mm^3^ (**: p=0,009; ***: p=0,0007; mean ± SD; two-way ANOVA, Bonferroni post-hoc test).

Taken together, our results show nuclear translocation of pSTAT3 and NF-kB in the striatum of STING ki mice, consistent with a robust activation of type I IFN and NF-kB/inflammasome dependent signaling in STING ki mice.

### Type I IFN and NF-kB/inflammasome signaling contribute to neuroinflammation in STING ki mice

We next sought to investigate the contribution of the type I IFN and NF-kB/inflammasome signaling pathways to neuroinflammation in STING ki mice. In order to test the involvement of the IFN dependent pathway, we crossed STING ki mice with mice deficient for type I IFN receptor (*Ifnar1^−/−^*); in order to test the involvement of the NF-kB/inflammasome pathway, we crossed STING ki mice with mice deficient for caspase-1 (*Casp1*^−/−^).

We first analyzed ISGs and NF-kB dependent gene expression by quantitative real time PCR in adult *Ifnar1^−/−^* and *Casp1*^−/−^ mice (Figure 6A-E). The N153S STING-induced increase in the interferon dependent genes *Ifi44 and Mx-1* was abrogated in *Ifnar1*^−/−^ mice (Figure 6A and B), but unaltered in *Casp1*^−/−^ mice. This is consistent with the dependence of *Ifi44 and Mx-1* on type I interferon signaling. The N153S STING-induced increase in expression of *Il-1*β (Figure 6C) was abrogated in *Casp1*^−/−^ mice, but not in *Ifnar1*^−/−^ mice, consistent with the fact that *Il-1*β is NF-κB dependent. The N153S STING-induced increase in *Ip-10* expression was 17-fold in WT (Figure 6D, p= 0,0030844), 12-fold in *Ifnar1^−/−^* (p= 0,002589) and 6-fold in *Casp 1*^−/−^ (p= 0,0041598), i.e. not statistically different between WT and *Ifnar1^−/−^* mice (p= 0,4398) or *Casp 1*^−/−^ mice (p= 0,0623). Expression of *Tnfa* (Figure 6E) was not different between STING ki and STING WT in this age group, consistent with Figure 5A, and unaltered in *Ifnar1^−/−^* and *Casp1*^−/−^ mice. Expression of *Sting1* was also not altered in any on the genotypes (figure supplement 2A), confirming that alteration in transgene expression did not underlie the observed effects. Collectively, these analyses confirm the activation of both IFN signaling and NF-κB/inflammasome signaling in the brain of STING ki mice. *Ifnar1^−/−^* selectively interferes with the interferon dependent pathway whereas *Casp1*^−/−^ interferes with the NF-kB/inflammasome pathway.

**Figure 6.**
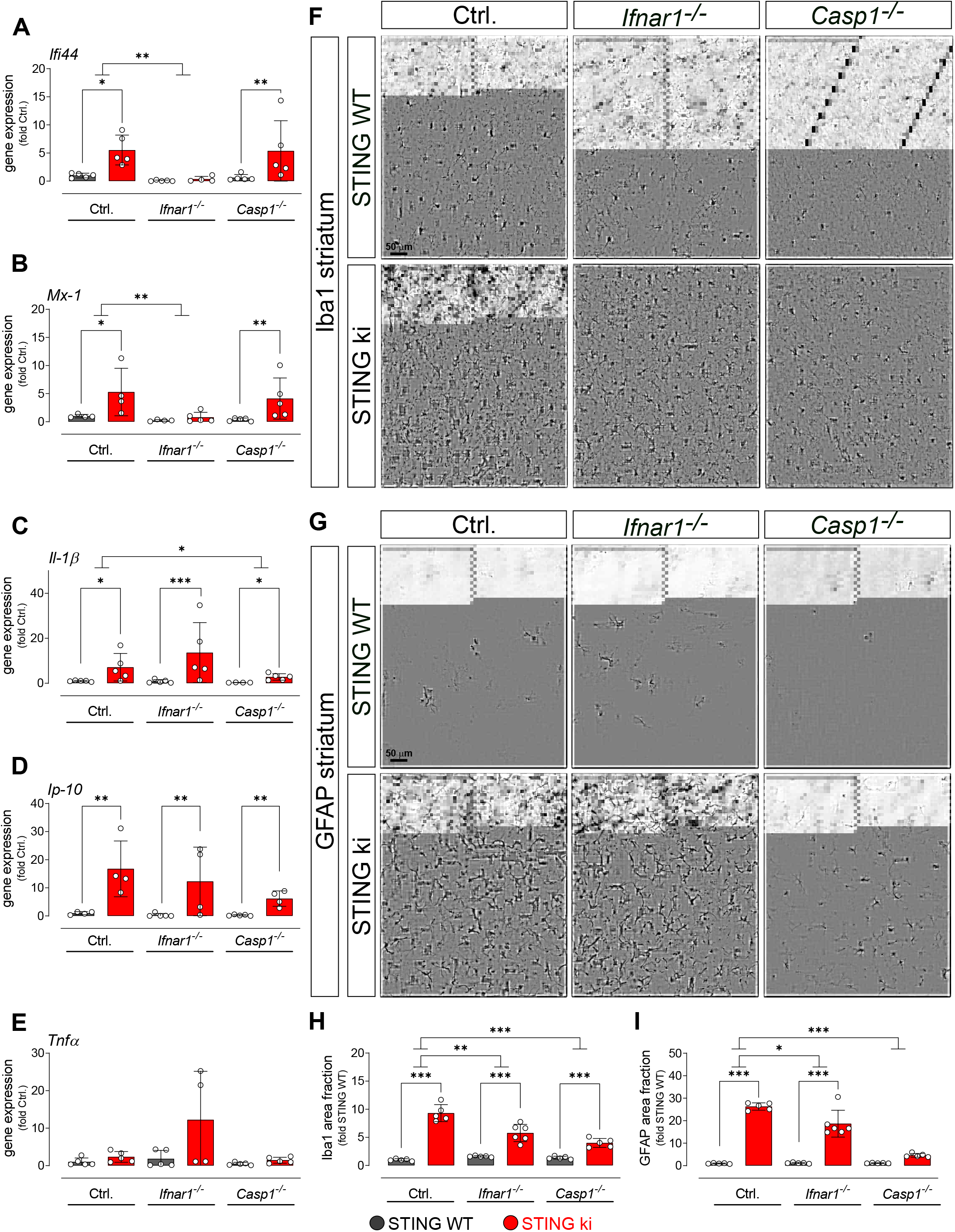
Neuroinflammation in adult double transgenic mice with STING ki and knock-out for Ifnar1 or Caspase-1. (A-E) Expression of IFN related genes and NF-κB/inflammasome related genes in the frontal cortex of adult STING WT and STING ki mice (ANOVA with Tukey HSD post-hoc test). (A) *Ifi44* (*: p= 0,01414; **: p= 0,0037655; for interaction between Ctrl. and and *Ifnar1^−/−^*, **: p= 0,005101). (B) *Mx-1* (*: p= 0,02823; **: p= 0,00573). (C) *Il-1b* (Ctrl. background *: p= 0,04534; **: p= 0, 005405; *Casp1^−/−^ background* *: p= 0,0107096; for interaction between Ctrl. and *Casp1^−/−^*, *: p= 0,01298). (D) *Ip-10* (Ctrl. background **: p= 0,0030844; *Ifnar1^−/−^* background: p= 0,025893; *Casp1^−/−^ background* **: p= 0,0041598). (E) *Tnfα* (all differences n.s.). *Sting1* expression is shown on figure supplement 2A. (F) Representative images of striatal sections stained for the microglia marker Iba1. Sections were obtained from adult STING WT (upper images) or STING ki (lower images) mice on a background of interferon a receptor knockout (*Ifnar1^−/−^*), caspase 1 knockout (*Casp1^−/−^*) or *Ifnar1^+/+^*, *Casp1^+/+^* (Ctrl.). Scale bar: 50 μm. (G) Representative images of striatal sections stained for the astroglia marker GFAP from STING WT (upper images) or STING ki (lower images) mice on a background of interferon a receptor knockout (*Ifnar1^−/−^*), caspase 1 knockout (*Casp1^−/−^*) or *Ifnar1^+/+^*, *Casp1^+/+^* (Ctrl.). Scale bar: 50 μm. (H) Area fraction positive for Iba1, normalized to the mean of STING WT brains (differences in ^+/+^ mice ***: p= 0,0000001; for *Ifnar1^−/−^* ***: p=0,000003; for *Casp1^−/−^* ***: p= 0,0029374; two-way ANOVA with Bonferroni post-hoc test). (I) Area fraction positive for GFAP, normalized to STING WT on Ctrl. Background (***: p= 0,0000 for STING WT vs STING ki on Ctrl.; ***: p= 0,0000 on *Ifnar1^−/−^*; background, ***: p=xyz on Casp1−/− background; two-way ANOVA with Bonferroni post-hoc test).

We next measured glial activation in *Ifnar1^−/−^* and *Casp1*^−/−^ mice. N153S STING-induced activation of microglia was still observed in adult *Ifnar1^−/−^* and *Casp1*^−/−^ mice (Figure 6F and H, fold increase and p values in legend). In order to determine whether the extent of N153S STING-induced microglia activation was significantly different between *Ifnar1^−/−^* mice and Ctrl., we calculated the interaction between the factors ‘STING genotype’ and ‘*Ifnar1* genotype’ in two-way ANOVA. The extent of microglia activation was significantly smaller in adult *Ifnar1^−/−^* mice (p= 0,004724) than in Ctrl. Similarly, the extent of N153S STING-induced microglia activation was significantly smaller in *Casp1*^−/−^ mice (p= 0,000021). The extent of N153S STING-induced microglia activation did not differ between *Ifnar1^−/−^* and *Casp1*^−/−^ (p= 0,0805731). Collectively, these findings suggest that N153S STING-induced microglia activation depends both on type 1 IFN signaling and on NF-kB/inflammasome signaling.

The extent of N153S STING-induced astroglia activation (Figure 6G and I) was significantly reduced from 26-fold in Ctrl. to 16-fold in *Ifnar1^−/−^* (p=0,0158378 for interaction of 2-way ANOVA) and to 4-fold in *Casp1*^−/−^ (p<0.00001 for interaction of 2-way ANOVA), suggesting that astroglia activation might depend more on NF-kB/inflammasome signaling than on interferon dependent signaling.

### Type I IFN and NF-kB/inflammasome signaling contribute to neurodegeneration in STING ki mice

Next, we asked whether the N153S STING-induced degeneration of dopaminergic axon terminals in the striatum is abrogated in adult *Ifnar1^−/−^* and *Casp 1*^−/−^ mice (Figure 7). Interestingly, the density of dopaminergic axon terminals in the striatum was already lower in *Ifnar1^−/−^* mice and *Casp 1*^−/−^ mice with WT STING than in control *Ifnar1^+/+^* and *Casp1*^+/+^ mice (p= 0,0143071 and p= 0,0000248, two-way ANOVA). These findings suggest that *Ifnar1* and *Casp1* are required for proper proliferation, maturation and/or maintenance of dopaminergic neurons and their axon terminals. Indeed, the Wnt-β-catenin pathway is regulated by interferons (Kovács et al., 2019) and regulates the differentiation of midbrain dopaminergic neurons (Szegő et al., 2017). Furthermore, IL-1β induces the differentiation of dopaminergic neurons (Ling et al., 1998; Rodriguez-Pallares et al., 2005), and we found marginally reduced *Il-1b* expression in adult *Casp1^−/−^* mice (Figure 6C STING WT).

**Figure 7.**
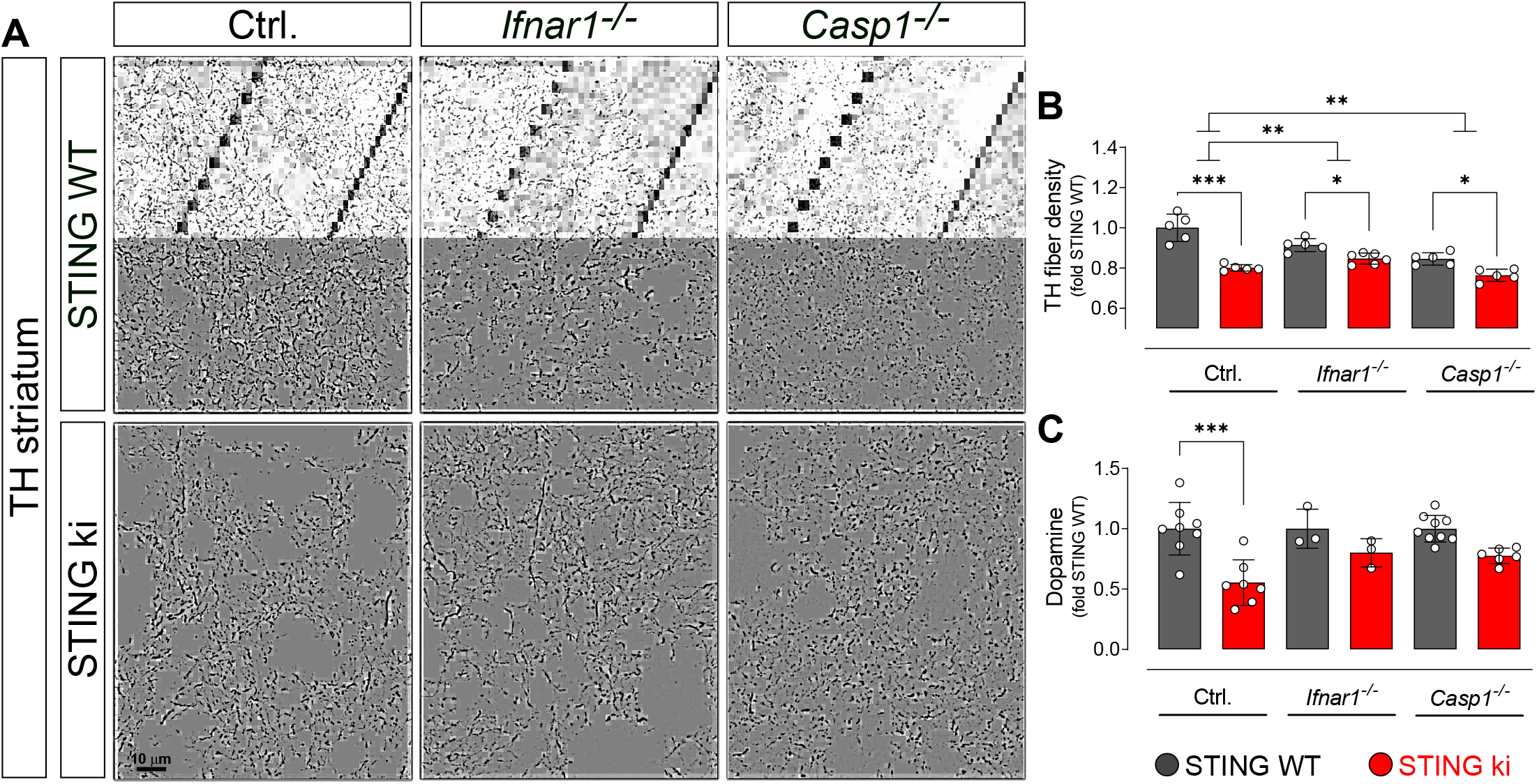
Degeneration of dopaminergic neurons in double transgenic mice with STING N153S/WT ki and knock-out for Ifnar1 or Caspase-1. (A) Representative images of striatal sections stained for tyrosine hydroxylase (TH) from STING WT (upper images) or STING ki (lower images) mice on a background of interferon a receptor knockout (*Ifnar1^−/−^*), caspase-1 knockout (*Casp1^−/−^*) or *Ifnar1^+/+^*, *Casp1^+/+^* (Ctrl.). Scale bar: 10 μm. (B) Area fraction positive for TH, normalized to STING WT on Ctrl. background (***: p=0,0000 for Ctrl. background; *: p= 0,043 for *Ifnar1^−/−^*; **: p=0,0126845 for *Casp1^−/−^*; for interaction between Ctrl. background and *Ifnar1^−/−^*: p= 0,00157; between Ctrl. and *Casp1^−/−^* : p= 0,007326; two-way ANOVA with Bonferroni post-hoc test). (C) Concentration of dopamine in striatal lysates of STING WT and STING ki mice, normalized to STING WT on Ctrl. background. (***: p= 0,0005; t-test). Dopamine metabolism is shown on figure supplement 2B.

Both in *Ifnar1^−/−^* and in *Casp 1*^−/−^ mice, the N153S STING-induced degeneration of dopaminergic axon terminals in the striatum was less pronounced than in the control *Ifnar1^+/+-^* and *Casp1*^+/+^ mice (p= 0,00157 and p= 0,007326 for interaction in two-way ANOVA). N153S STING-induced fiber loss was not statistically different between *Ifnar1^−/−^* and *Casp1*^−/−^ mice (p= 0,468 for interaction). Consistent with the less pronounced degeneration of dopaminergic axon terminals, the N153S STING-induced reduction in striatal dopamine and increase in its metabolism was not observed in *Ifnar1^−/−^* and *Casp1*^−/−^ mice (Figure 7C, figure supplement 2B). These findings suggest that both pathways contribute to the degeneration of dopaminergic axon terminals, and blocking either pathway could reduce neurodegeneration.

## Discussion

In this work, we demonstrated that the expression of the constitutively active STING variant N153S causes neuroinflammation, followed by degeneration of dopaminergic neurons and aSyn pathology. N153S STING-induced microglia activation and degeneration of dopaminergic neurons involve both type I IFN-dependent and NF-κB/inflammasome-dependend signaling, while astroglia activation might depend predominantly on the NF-κB/inflammasome pathway.

### Constitutive STING activation causes neuroinflammation

STING is expressed in the central nervous system. Its expression is highest in microglia, but it can also be detected in astroglia and neurons (Jeffries and Marriott, 2017). Microglia are the primary immune cells of the central nervous system (Wolf et al., 2017) and highly activated upon pathogen invasion or tissue damage. Microglia activation in STING ki mice (Figure 1 and 2) is therefore in line with the increased inflammatory phenotype found in the lungs and spleen of STING ki mice (Luksch et al., 2019; Siedel et al., 2020) and in patients with STING-associated vasculopathy with onset in infancy syndrome (Liu et al. 2014).

Consistent with this morphologically defined neuroinflammatory phenotype, we observed increased expression of the ISGs *Ifi44* and *Mx-1* (Figure 4). These effects are associated with interferon signaling, the main pathway downstream of STING (Decout et al., 2021), demonstrating that expressing the N153S mutant of STING indeed activates interferon dependent pathways. Microglia activation was reduced when N153S STING was expressed in mice deficient for Ifnar1 (*Ifnar1*^−/−^, Figure 6H), confirming that the type I IFN pathway is involved in STING-dependent microglia activation as previously demonstrated by others (Warner et al., 2017). However, microglia activation by N153S STING was not completely blocked in *Ifnar1^−/−^* mice, suggesting that additional pathways are involved.

Indeed, next to type I IFN signaling, we also observed increased expression of *Il-1b* and *Tnfa* (Figure 4), confirming the activation of the NF-κB/inflammasome pathway in STING ki mice (Balka et al., 2020; Balka and De Nardo, 2021; B. C. Liu et al., 2018). Consistently, in STING ki mice, nuclear translocation of NF-κB was observed significantly more often (Figure 5). We also observed increased expression of *Casp1*, indicating that the inflammasome pathways are activated by constitutively active STING, consistent with previous work about systemic inflammation (Luksch et al., 2019). Accordingly, the extent of microglia activation by N153S STING was reduced in *Casp1^−/−^* mice (Figure 6H). Therefore, our results suggest that STING-dependent microglia activation involves inflammasome activation, in addition to type I IFN-dependent pathways.

Next to the activation of microglia, we also observed activation of astroglia (Figures 1 and 2). STING-induced astroglia activation was only blunted in *Ifnar1^−/−^* mice, but it was completely blocked in *Casp1^−/−^* mice (Figure 6I). This finding suggests that chronic STING-induced astroglia activation might depend mainly on NF-κB/inflammasome signaling. Astroglia activation can result directly from the expression of N153S STING in astroglia and indirectly through the activation of microglia (Kwon and Koh, 2020; Wolf et al., 2017). The differential effect in *Ifnar1^−/−^* and *Casp1^−/−^* mice suggests that astroglia activation is not only a downstream consequence of microglia activation.

### Chronic STING activation leads to the degeneration of dopaminergic neurons

We observed a reduced number of TH-positive neurons in the *substantia nigra* of STING ki mice, a reduced density of dopaminergic axon terminals and a reduced concentration of striatal dopamine (Figure 1G-I). These changes occurred in adult mice and were not present in juvenile mice (Figure 2G-I). The degeneration of dopaminergic neurons thus is a consequence of the inflammatory changes in our model.

On the *Ifnar1^−/−^* and *Casp1^−/−^* backgrounds, the effect of STING ki was difficult to assess. Yet, the relative reduction in TH fiber density in STING ki mice, as compared to STING WT mice, was smaller on the *Ifnar1^−/−^* and *Casp1^−/−^* backgrounds than in controls (Figure 7B), as was the extent of dopamine depletion F (Figure 7C). These findings are consistent with the partial rescue of microglia activation in *Ifnar1^−/−^* and *Casp1^−/−^* mice (Figure 6F and H). They suggest that both pathways contribute to the degeneration of dopaminergic neurons. In further studies, the developmental effects of *Ifnar1* and *Casp1* deficiency could be circumvented by using conditional knockout mice or pharmacological inhibitors.

In our model, degeneration of dopaminergic neurons and their axon terminals in STING ki mice is likely a secondary effect of glia activation and secretion of inflammatory cytokines – in line with previous findings demonstrating a role for inflammation in the pathogenesis of PD (Hall et al., 2018; Mollenhauer et al., 2019). In addition, constant STING activity within dopaminergic neurons could contribute to their degeneration – both in our STING ki mice and in the pathogenesis of PD. Indeed, dopaminergic neurons accumulate oxidative damage as a consequence of dopamine synthesis and electrical pacemaking activity (Guzman et al., 2010), and even moderate oxidative stress can stimulate the STING pathway (Sliter et al., 2018; West et al., 2015). Further work is required to determine the importance of STING activation in dopaminergic neurons for their degeneration in this model.

### Accumulation of protein aggregates in STING ki mice

We observed an accumulation of pathological aSyn and an increased number of Thioflavin S positive cells in STING ki mice (Figure 3A-E). These findings suggest that chronic STING activation induces aSyn pathology. They are consistent with the recent observation that priming rats with a mimic of viral dsDNA precipitates aSyn pathology (Olsen et al., 2019) and with the aSyn aggregation following viral encephalitis (Bantle et al., 2021). Inflammatory signals are therefore active promotors of aSyn pathology and not only responsive to aSyn pathology.

The mechanism by which prolonged neuroinflammation leads to aSyn pathology is still unknown. Both inflammation and IFN induce the expression of the double-stranded (ds) RNA-dependent protein kinase (PKR) (Gal-Ben-Ari et al., 2019). PKR can phosphorylate aSyn at serine 129, resulting in aSyn pathology (Reimer et al., 2018). Moreover, clearance of aSyn aggregates occurs primarily through autophagy (Ebrahimi-Fakhari et al., 2011), and INFα increases expression of mammalian target of rapamycin (mTOR) (Liu et al., 2016), which is expected to reduce autophagy initiation. On the other hand, acute STING activation can induce autophagy (Hopfner and Hornung, 2020; Y. Liu et al., 2018; Moretti et al., 2017). aSyn pathology in our model therefore could be explained by an exhaustion of the autophagy machinery, as it was suggested recently (Bido et al., 2021), and the overall effect of inflammatory pathways on autophagy and aSyn pathology could be bimodal. Accordingly, degeneration of dopaminergic neurons and accumulation of aSyn aggregates was also observed in mice deficient for IFNβ (Ejlerskov et al., 2015; Magalhaes et al., 2021).

## Conclusions

In this work, we observed the degeneration of dopaminergic neurons in mice expressing a constitutively active STING variant. These findings support the hypothesis that chronic neuroinflammation is sufficient to trigger degeneration of dopaminergic neurons and aSyn pathology. They indicate that neuroinflammation plays an active role in the pathogenesis of PD and could be a promising therapeutic target. Further work is required to determine the precise signaling pathway by which STING activation causes aSyn pathology and dopamine neuron degeneration.

## Declarations

### Ethical Approval and Consent to participate

Animals were approved by local authorities (Landesdirektion Dresden)and conducted in accordance with guidelines of the Federation for European Laboratory Animal Science Associations (FELASA).

### Consent for publication

All authors approved the manuscript.

### Availability of data and materials

The dataset supporting the conclusions of this article is included within the article and its additional files.

### Competing interests

The authors declare that they have no competing interests.

### Funding

This work was supported by the German Research Foundation (DFG) SFB TRR237, B18 and FA-658-3-1.

### Authors’ contributions

EMSz, ARW, BHF and HL conceived research. EMSz, LM, NB and HL performed research and analyzed data. EMSz, ARW, BHF and HL wrote the first draft of the manuscript. All authors contributed and approved the manuscript.

## Acknowledgements

We thank Min Ae Lee-Kirsch (TU Dresden) for very helpful comments on the manuscript and Andrea Kempe, Annett Böhme, Katrin Höhne and Kristin Wogan for excellent technical assistance.

## List of Abbreviations

aSyn: alpha-synuclein
Casp1: Caspase1
cGAS: cyclic GMP-AMP synthase
dsDNA: double strand DNA
GFAP: glial fibrillary acidic protein
Iba1: ionized calcium-binding adapter molecule 1
Ifi44: interferon induced protein 44
IFN: interferon
Ifnar1: interferon alpha receptor1
Il-1β: interleukin 1 beta
Ip-10: interferon-gamma induced protein 10 kD
IRF3: interferon regulatory factor 3
ISG: interferon-stimulated gene
ki: knock in
Mx1: interferon-induced GTP-binding protein
NF-kB: nuclear factor ’kappa-light-chain-enhancer’ of activated B-cells
NRLP3: the nucleotide-binding oligomerization domain (NOD), leucine-rich repeat (LRR)-containing protein 3
PD: Parkinson’s disease
SAVI: STING-associated vasculopathy with onset in infancy
SN: substantia nigra
STAT3: signal transducer and activator of transcription 3
STING: stimulator of interferon genes
TH: tyrosine hydroxylase
TNFβ: tumor necrosis factor beta
WT: wild type

## Supplemental Information

**Figure supplement 1 related to Figures 1 and 2.**
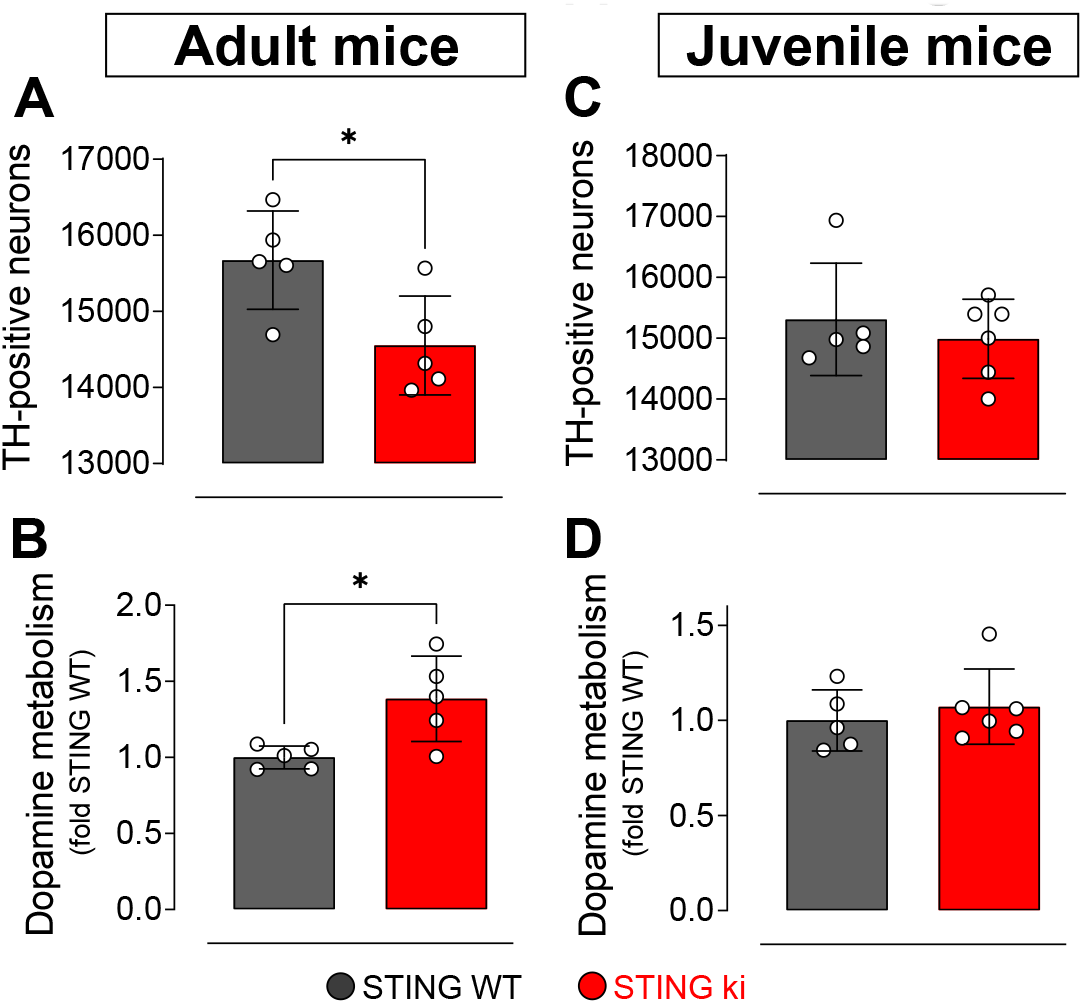
Number of TH-positive neurons in the substantia nigra and dopamine metabolism in the striatum of STING WT and STING ki mice. (A) Number of TH-positive neurons in the substantia nigra of adult mice (p=0,0257; t-test). (B) Dopamine metabolism (concentration of dopamine metabolites DOPAC + HVA) / dopamine in adult mice (p=0,0179; t-test). (C) Number of TH-positive neurons in the substantia nigra of juvenile mice (p=0,5188; t-test). (D) Dopamine metabolism in juvenile mice (p=0,9545; t-test).

**Figure supplement 2 related to Figures 6 and 7.**
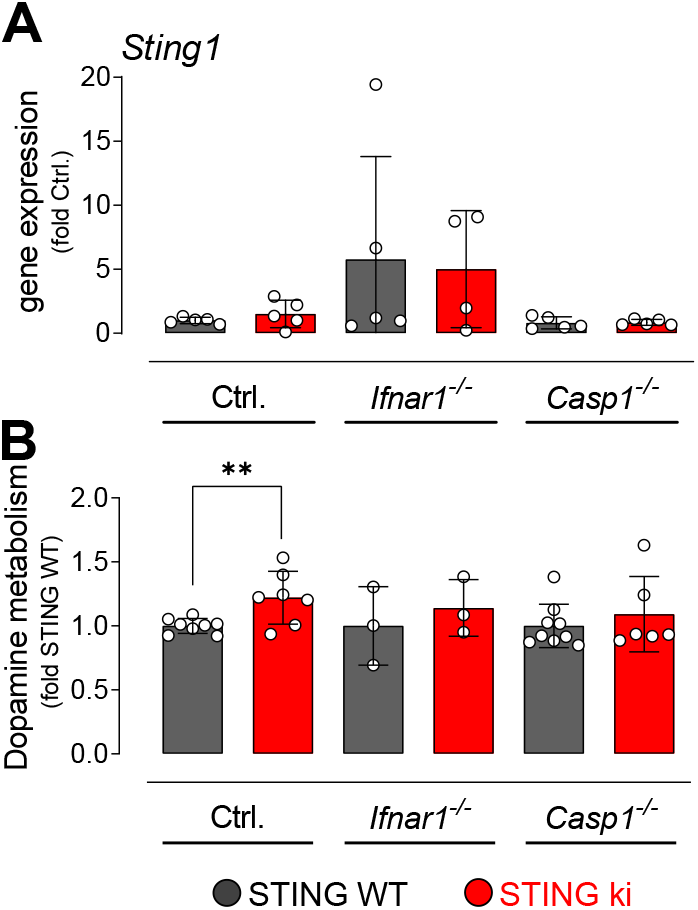
*Sting1* expression and dopamine metabolism in KO animals. (A) Expression of *Sting1* in the frontal cortex of STING WT and STING ki mice on a background of interferon a receptor knockout (*Ifnar1^−/−^*), caspase 1 knockout (*Casp1^−/−^*) or *Ifnar1^+/+^*, *Casp1^+/+^* (Ctrl.). (all differences n.s., 2-way ANOVA). (B) Dopamine metabolism in STING WT and STING ki mice. (**: p= 0,0073; 2-way ANOVA).

**Supplemental Table S1.**
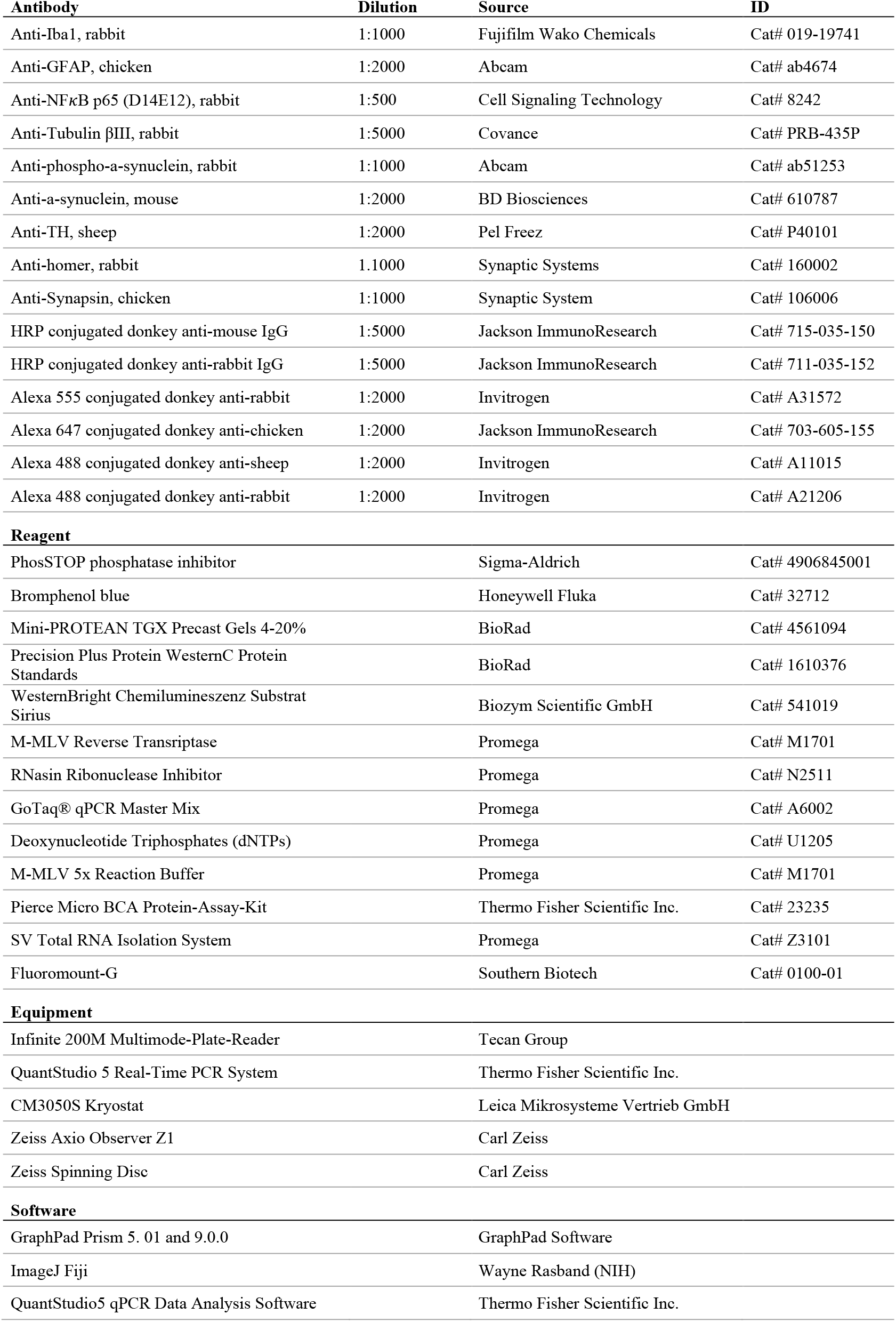
List of materials and antibody dilutions

| **Antibody** | **Dilution** | **Source** | **ID** |
| --- | --- | --- | --- |
| Anti-Iba1, rabbit | 1:1000 | Fujifilm Wako Chemicals | Cat# 019-19741 |
| Anti-GFAP, chicken | 1:2000 | Abcam | Cat# ab4674 |
| Anti-NFκB p65 (D14E12), rabbit | 1:500 | Cell Signaling Technology | Cat# 8242 |
| Anti-Tubulin βIII, rabbit | 1:5000 | Covance | Cat# PRB-435P |
| Anti-phospho-a-synuclein, rabbit | 1:1000 | Abcam | Cat# ab51253 |
| Anti-a-synuclein, mouse | 1:2000 | BD Biosciences | Cat# 610787 |
| Anti-TH, sheep | 1:2000 | Pel Freez | Cat# P40101 |
| Anti-homer, rabbit | 1:1000 | Synaptic Systems | Cat# 160002 |
| Anti-Synapsin, chicken | 1:1000 | Synaptic System | Cat# 106006 |
| HRP conjugated donkey anti-mouse IgG | 1:5000 | Jackson ImmunoResearch | Cat# 715-035-150 |
| HRP conjugated donkey anti-rabbit IgG | 1:5000 | Jackson ImmunoResearch | Cat# 711-035-152 |
| Alexa 555 conjugated donkey anti-rabbit | 1:2000 | Invitrogen | Cat# A31572 |
| Alexa 647 conjugated donkey anti-chicken | 1:2000 | Jackson ImmunoResearch | Cat# 703-605-155 |
| Alexa 488 conjugated donkey anti-sheep | 1:2000 | Invitrogen | Cat# A11015 |
| Alexa 488 conjugated donkey anti-rabbit | 1:2000 | Invitrogen | Cat# A21206 |
| **Reagent** |  |  |  |
| PhosSTOP phosphatase inhibitor |  | Sigma-Aldrich | Cat# 4906845001 |
| Bromphenol blue |  | Honeywell Fluka | Cat# 32712 |
| Mini-PROTEAN TGX Precast Gels 4-20% |  | BioRad | Cat# 4561094 |
| Precision Plus Protein WesternC Protein Standards |  | BioRad | Cat# 1610376 |
| WesternBright Chemilumineszenz Substrat Sirius |  | Biozym Scientific GmbH | Cat# 541019 |
| M-MLV Reverse Transcriptase |  | Promega | Cat# M1701 |
| RNasin Ribonuclease Inhibitor |  | Promega | Cat# N2511 |
| GoTaq® qPCR Master Mix |  | Promega | Cat# A6002 |
| Deoxynucleotide Triphosphates (dNTPs) |  | Promega | Cat# U1205 |
| M-MLV 5x Reaction Buffer |  | Promega | Cat# M1701 |
| Pierce Micro BCA Protein-Assay-Kit |  | Thermo Fisher Scientific Inc. | Cat# 23235 |
| SV Total RNA Isolation System |  | Promega | Cat# Z3101 |
| Fluoromount-G |  | Southern Biotech | Cat# 0100-01 |
| **Equipment** |  |  |  |
| Infinite 200M Multimode-Plate-Reader |  | Tecan Group |  |
| QuantStudio 5 Real-Time PCR System |  | Thermo Fisher Scientific Inc. |  |
| CM3050S Kryostat |  | Leica Mikrosysteme Vertrieb GmbH |  |
| Zeiss Axio Observer Z1 |  | Carl Zeiss |  |
| Zeiss Spinning Disc |  | Carl Zeiss |  |
| **Software** |  |  |  |
| GraphPad Prism 5. 01 and 9.0.0 |  | GraphPad Software |  |
| ImageJ Fiji |  | Wayne Rasband (NIH) |  |
| QuantStudio5 qPCR Data Analysis Software |  | Thermo Fisher Scientific Inc. |  |

**Supplemental Table S2.**
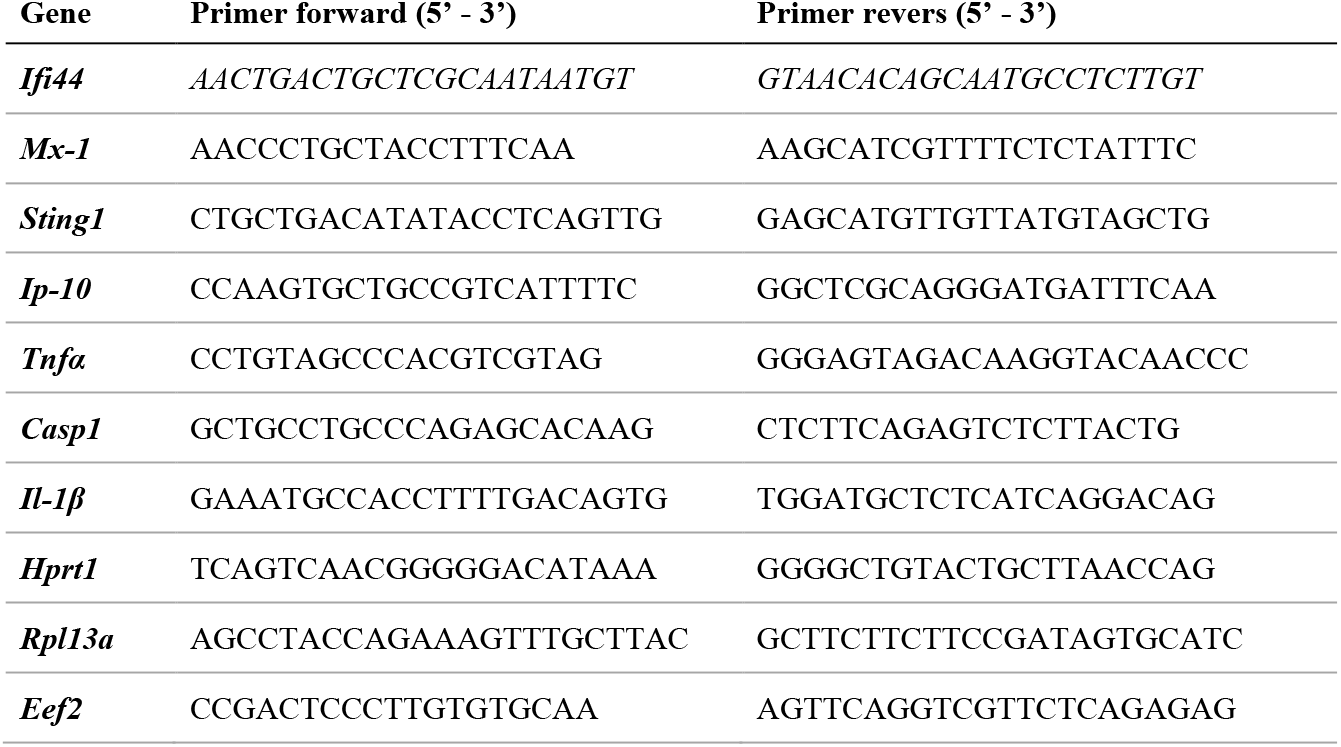
List of qRT PCR Primers

**Supplemental Table S4.** Sample numbers in each analysis.

| Assay | Group -- number of animals used/excluded |  |  |  |  |  | Reason* |
| --- | --- | --- | --- | --- | --- | --- | --- |
| Western blot | STING WT adult<br>5/0 | STING ki adult<br>5/0 |  |  |  |  |  |
| qPCR | STING WT adult | STING ki adult | STING WT juvenile | STING ki juvenile |  |  |  |
| *Ifi44* | 5/0 | 5/0 | 5/0 | 5/0 |  |  |  |
| *Ip-10* | 4/1 | 4/1 | 5/0 | 5/0 |  |  | Outlier |
| *Mx-1* | 4/1 | 4/1 | 5/0 | 5/0 |  |  | Outlier |
| *Tnfa1* | 5/0 | 5/0 | 5/0 | 5/0 |  |  |  |
| *Casp1* | 5/0 | 4/1 | 5/0 | 5/0 |  |  | Outlier |
| *Il-1b* | 4/1 | 5/0 | 4/1 | 5/0 |  |  | Outlier |
| *Sting1* | 5/0 | 5/0 | 5/0 | 4/1 |  |  | Outlier |
|  | STING WT | STING ki | STING WT *Ifnar1<sup>-/-</sup>* | STING ki *Ifnar1<sup>-/-</sup>* | STING WT *Casp1<sup>-/-</sup>* | STING ki *Casp1<sup>-/-</sup>* |  |
| *Ifi44* | 5/0 | 5/0 | 5/0 | 5/0 | 4/1 | 5/0 | Outlier |
| *Ip-10* | 4/1 | 4/1 | 4/1 | 4/1 | 4/1 | 4/1 | Outlier |
| *Mx-1* | 4/1 | 4/1 | 4/1 | 5/0 | 5/0 | 5/0 | Outlier |
| *Tnfa1* | 5/0 | 5/0 | 5/0 | 4/1 | 5/0 | 5/0 | Outlier |
| *Il6* | 4/1 | 5/0 | 5/0 | 4/1 | 4/1 | 4/1 | Outlier |
| *Casp1* | 5/0 | 5/0 | 5/0 | 5/0 | 5/0 | 5/0 | Outlier |
| *Il-1b* | 5/0 | 5/0 | 4/1 | 5/0 | 4/1 | 5/0 | Outlier |
| Histology | STING WT adult<br>5/0 | STING ki adult<br>5/0 | STING WT juvenile<br>5/0 | STING ki juvenile<br>6/0 |  |  |  |
|  | STING WT | STING ki | STING WT, *Ifnar1<sup>-/-</sup>* | STING ki, *Ifnar1<sup>-/-</sup>* | STING WT, *Casp1<sup>-/-</sup>* | STING ki, *Casp1<sup>-/-</sup>* |  |
|  | 5/0 | 5/0 | 5/0 | 6/0 | 5/0 | 5/0 |  |
| HPLC | STING WT adult<br>5/0 | STING ki adult<br>5/0 | STING WT juvenile<br>5/0 | STING ki juvenile<br>5/0 |  |  |  |
|  | STING WT | STING ki | STING WT, *Ifnar1<sup>-/-</sup>* | STING ki, *Ifnar1<sup>-/-</sup>* | STING WT, *Casp1<sup>-/-</sup>* | STING ki, *Casp1<sup>-/-</sup>* |  |
|  | 8/0 | 7/0 | 3/0 | 3/0 | 9/0 | 6/0 |  |

\*Grubbs`test was used to detect outliers (Prism).

## Notes

### Competing Interest Statement

The authors have declared no competing interest.

### Summary of Updates

updated "author contributions" statement.

